# Distribution of multi-unit pitch responses recorded intracranially from human auditory cortex

**DOI:** 10.1101/2021.10.22.465330

**Authors:** Joel I Berger, Phillip E Gander, Yukiko Kikuchi, Sukhbinder Kumar, Christopher Kovach, Hiroyuki Oya, Hiroto Kawasaki, Matthew A Howard, Timothy D Griffiths

**Author notes:** **Corresponding author:** Joel I Berger. **Competing Interest Statement:** The authors declare no competing financial interests.

## Abstract

The perception of pitch requires the abstraction of stimulus properties related to the spectrotemporal structure of sound. Previous studies utilizing both animal electrophysiology and human imaging have indicated the presence of a center for pitch representation in the auditory cortex. Recent data from our own group - examining local field potentials (LFPs) in humans - indicate more widely distributed pitch-associated responses within the auditory cortex (Gander et al., 2019). To probe this with greater spatial resolution, we examined multi-unit activity related to three different auditory stimuli, in seven epilepsy patients who were implanted with high-impedance electrodes in auditory cortex for the clinical purpose of localizing seizures. The stimuli were regular-interval noise (RIN) with a pitch strength that is related to the temporal regularity, and pitch value determined by repetition rate, and harmonic complexes with missing fundamentals. We demonstrated increases in spiking activity in 69 of 104 (66%) responsive multiunit activity in auditory cortex due to pitch-associated stimuli. Importantly, these responses were distributed across the entire extent of Heschl’s gyrus (HG), in both primary and non-primary areas, rather than isolated to a specific region, and this finding was evident regardless of the stimulus presented. These findings are the first multi-unit pitch responses recorded from humans, and align with a recent study in macaques (Kikuchi et al., 2019) demonstrating that both local field potential and unit responses to pitch-inducing stimuli are distributed throughout auditory cortex.

**Significance Statement:** The perception of pitch is a fundamental acoustic attribute that is mediated by the auditory system. Despite its importance, there is still debate as to the precise areas responsible for its encoding, which may be due to differences in the recording measures and choices of stimuli used in previous studies. Here, we present the first study to measure multi-unit pitch responses in the auditory cortices of intracranially-implanted humans. Importantly, we demonstrate reliable responses to three different pitch-inducing paradigms that are distributed throughout Heschl’s gyrus, rather than being localized to a particular region. These data provide a bridge across animal and human studies, and aid in our understanding of the processing of a critical attribute of acoustic stimuli.

## Introduction

Pitch is a fundamental percept that is a critical aspect of music and voice perception, and sound segregation. Previous studies of the neural basis for pitch perception employed different categories of pitch-inducing stimuli. These include regular interval noise (RIN): noise that is iteratively delayed by a fixed time interval and added to the undelayed noise, resulting in a temporal regularity which increases the perceived salience of pitch as the number of iterations increases ([3]; [4]; [5]). This stimulus allows control of the long-term spectrum of the sound. Other studies have used harmonic complexes. Harmonics in the range one to ten are resolved by the cochlea, and are associated with strong pitch, while harmonics above ten contribute a less salient pitch ([6]; [7]). A variety of other pitch stimuli have also been used including Huggins pitch produced by creating a phase difference between the ears for a particular noise band (e.g. [8]). A neural mechanism that represents the pitch percept should represent that percept irrespective of the sensory stimulus from which the pitch is abstracted, and should only occur in regions of stimulus space associated with pitch: in terms of pitch value, above the lower limit of pitch (about 30 Hz in human; [9]).

Human functional imaging studies based on fMRI to measure blood-oxygenation-level dependent (BOLD) responses as an index of ensemble activity have compared pitch-associated stimuli with control stimuli ([10]; [11]; [12]; [13]; [8]). These studies have suggested a region in lateral HG and adjacent areas that shows greater activation to pitch, although not all studies support the idea ([14]; [15]).

The neurophysiological basis for pitch has been addressed in non-human primates (NHPs: Bendor and Wang, [16]). Employing strict criteria for the determination of pitch-sensitive single neurons in marmosets, Bendor and Wang demonstrated selectivity of responses in a region overlapping anterolateral A1 and belt regions of auditory cortex, consistent with the idea of pitch only being processed in a selective area of auditory cortex. However, recent NHP and human studies in which pitch responsiveness was defined based on responses to different pitch-associated stimuli with values above the lower limit of pitch have shown more widespread pitch-associated responses. These studies have been based on single-unit, multi-unit activity (MUA) and LFP responses in macaques ([2]) and LFPs in humans ([17]; [1]).

In this study we address the neural basis at the level of MUA recorded from human auditory cortex using two types of pitch-associated stimuli. This is the first human study to examine directly neuronal spiking activity in response to pitch-associated stimuli, as opposed to local field potentials or BOLD activity. It is important to examine at this level as most human studies at present utilize fMRI, with only a few examining LFPs, and BOLD activity may not correlate well with MUA ([18]). Moreover, MUA allows for a more direct comparison with animal studies of pitch, all of which use spiking activity as a primary response measure. The recordings from human primary and nonprimary areas allow us to test the hypothesis that responses associated with the pitch percept are localized in one cortical area. We also wished to examine whether different pitch-associated stimuli resulted in different spatial specificity, as a possible basis for the lack of concordance between the various studies. We employed RIN stimuli - varying either the pitch value or the pitch salience – and harmonic complex tones. Criteria for pitch responsiveness in the current study were increased spiking responses to the pitch-eliciting stimuli compared to control noise only occurring above the lower limit of pitch (~30 Hz in humans; [9]), and increasing spiking responses as pitch salience increased.

## Results

Figure 1 shows diagrams of these six conditions for the RIN iterations paradigm, wherein the number of delay-and-add cycles increases pitch salience. We also recorded responses to two other stimuli: RIN delays, in which the periodicity of the RIN is varied, to assess the preferred rate of MUA; Harmonic complex, where the stimuli produce a percept of either a 200 Hz or 500 Hz missing fundamental (see *Materials and Methods* for further details). The total percentage of pitch and non-pitch MUA across all subjects for the iterations, delays and harmonic complex paradigms resulted in 26/42 (61.90%), 29/42 (69.05%) and 14/20 (70.00%) of MUA showing pitch responses, respectively. Differences in the proportion of pitch/non-pitch MUA were tested with multiple Fisher’s exact tests. There was no significant difference between the proportion of units classified as pitch responsive for the harmonic complex paradigm compared to the iterations and delays paradigms, (*p* = 0.58 and *p* = 1.00 respectively, two-tailed Fisher’s exact tests). There was also no significant difference between the iterations and delays paradigms (*p* = 0.65).

**Figure 1.**
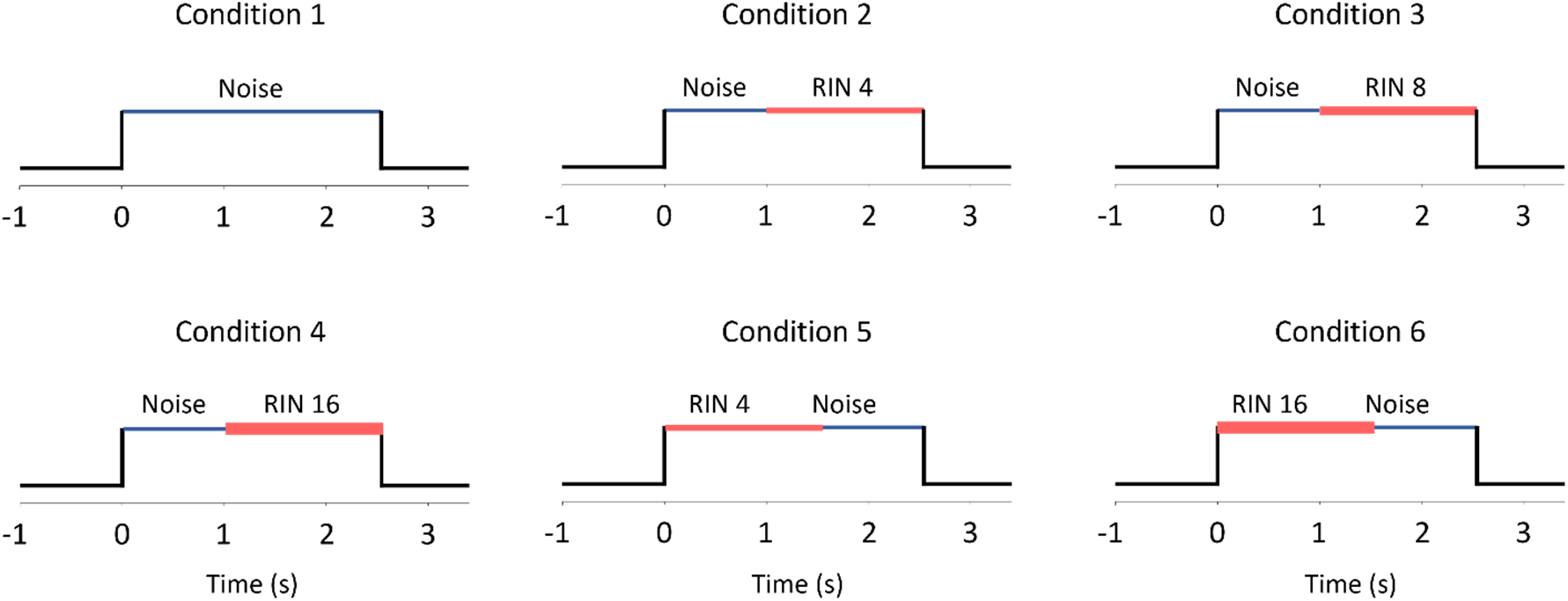
Diagrams of the stimuli used in the RIN iterations paradigm. Each subplot indicates a separate condition. RIN values show the number of delay-and-add cycles used to generate the stimuli, which results in a greater pitch percept as this number increases. Line thickness changes according to RIN value. Each trial consisted of 1 second of noise and 1.5 seconds of RIN.

### RIN iterations

Before examining the spatial distribution of pitch responses, it is useful to first examine the response patterns. Each panel in Figure 2 represents the grand mean normalized PSTH of all pitch-responsive MUA to each condition of the RIN iterations paradigm (*n* = 26 MUA, 50 trials per condition per MUA), wherein the RIN stimulus had a fixed periodicity of either 128 Hz or 256 Hz and only the number of iterations was incremented. On average, there was an initial onset response evident to the onset of the initial noise stimulus (time 0 in the first four panels, peak response between 138 – 151 ms post-stimulus) and a final offset response to all the stimuli. When no iterations were added, there was generally only an onset and then a sustained response to the ongoing noise, followed by a fast offset response (peak at 58 ms post-stimulus offset; Figure 2A). The peak response to the onset of the noise was between 128 and 156 ms when the noise was preceded by a pitch-like stimulus (Figures 2E and 2F). In five of the six conditions, noise was iterated with a delay-and-add algorithm to create the pitch-like percept (RIN). The peak onset responses to these RIN stimuli were between 111 and 114 ms when preceded by noise and did not systematically differ according to the salience of the pitch percept (Figures 2B, 2C and 2D). When preceded by silence, the peak of the onset response occurred between 108 and 122 ms (Figures 2E and 2F). In all these RIN conditions, a fast offset response was also present when the stimulus was followed by silence, with the peak occurring between 49 and 58 ms following the cessation of any auditory stimulation. The onset responses to the RIN stimuli were larger than the onset responses to the noise when both types of stimuli were preceded by silence, with a peak Δ firing rate of 15.35 Hz for noise, 20.86 Hz for 4-RIN and 23.39 Hz for 16-RIN. For the 16-RIN condition preceded by silence (i.e. the most responsive condition), mean baseline firing rates for grand mean MUA were 20.46 Hz (± 1.15 s.d.). Mean firing rates within 250 ms of stimulus onsets for this condition were 34.85 Hz (± 6.62 s.d.) for RIN and 28.65 Hz (± 2.33 s.d.) for noise.

**Figure 2.**
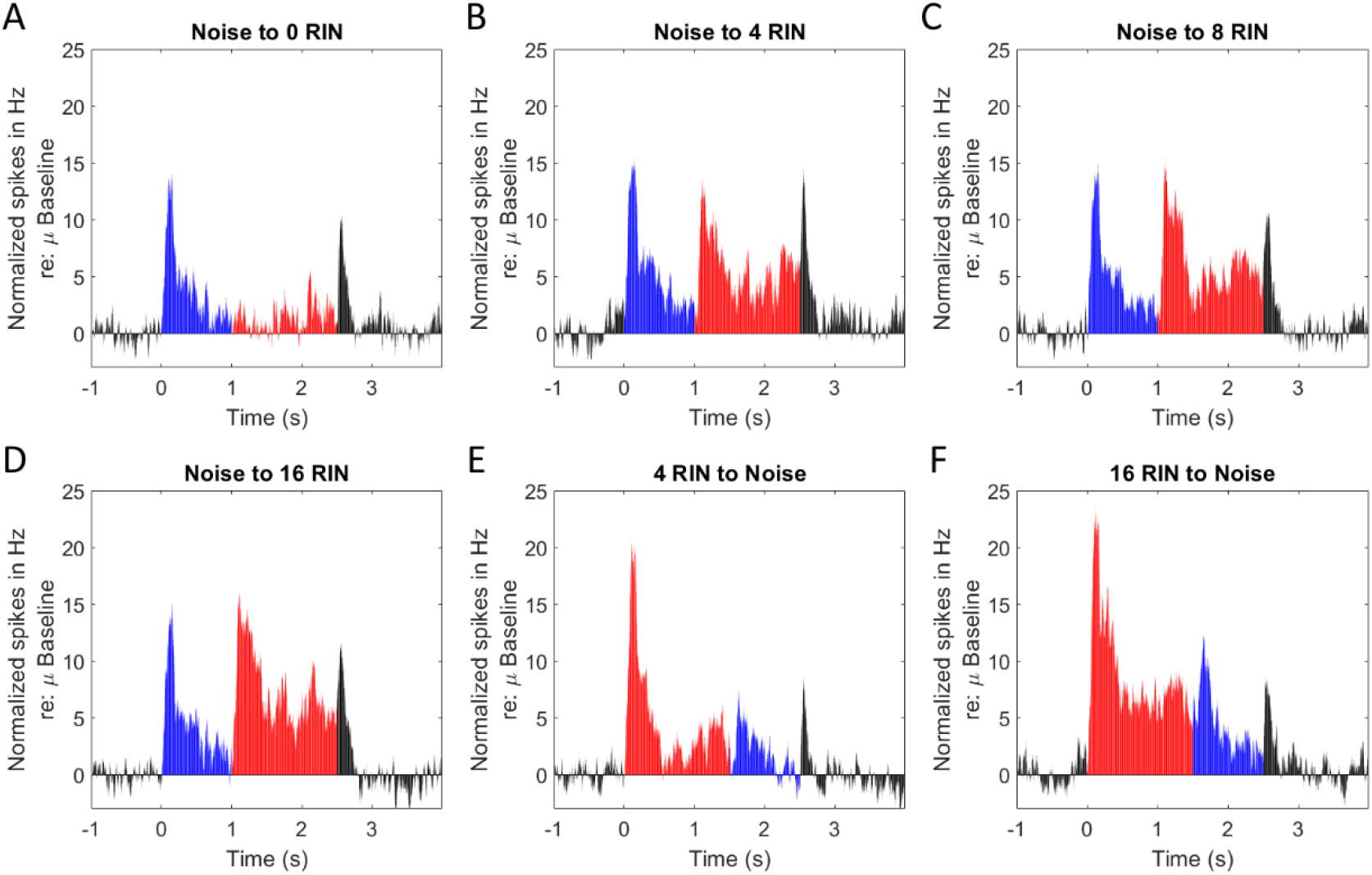
Grand mean PSTHs across all pitch-responsive MUA (1ms bins smoothed with a 25 ms window size moving average for display purposes), recorded in response to the RIN iterations paradigms (*n* = 26). Grand mean values shown are normalized (via subtraction prior to smoothing) relative to the mean firing during the pre-stimulus baseline and each subplot represents a different condition. Data are averaged across 128 Hz and 256 Hz stimuli. Red bars - RIN period; blue bars - noise period; black bars - silence period.

### RIN delays

Figure 3A shows example MUA PSTHs to the six different conditions of the RIN delays paradigm (50 trials per condition), and the corresponding trial raster is shown in Figure 3B. A total of 29 MUA (out of 42) were determined to be pitch responsive for these stimuli. Different MUA showed different best rates – in the current figure, this MUA responded preferentially to the 64Hz periodicity, with a peak Δ of 84.29 Hz relative to baseline; the normalized response to the onset of noise for this same condition was 30.69 Hz. The number of MUA with best rates for each of the frequencies above the lower limit of pitch were 2 for 32 Hz, 8 for 64 Hz, 3 for 128 Hz and 16 for 256 Hz.

**Figure 3.**
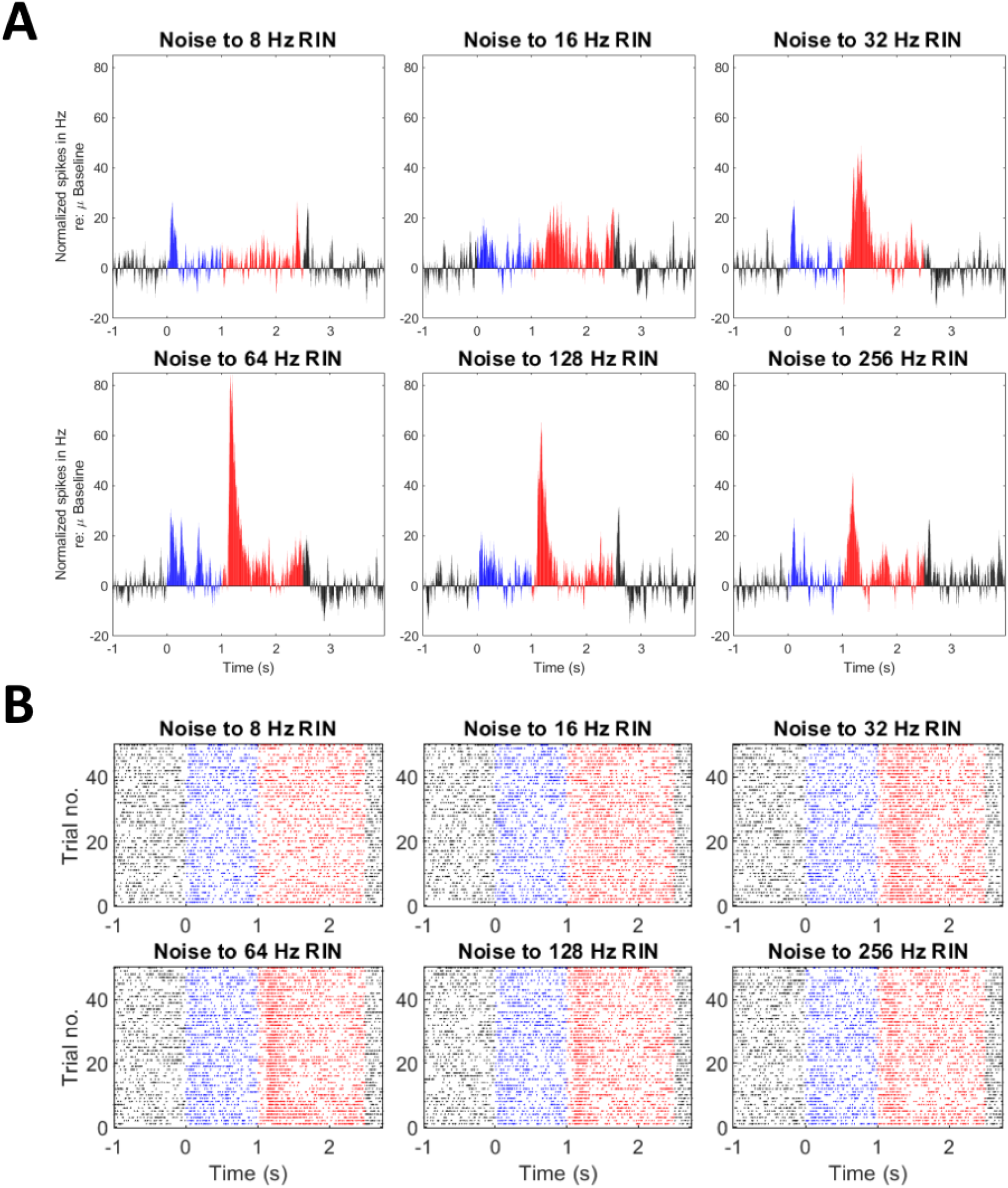
An example of normalized MUA PSTHs to the RIN delays paradigm (A), with corresponding spike time raster plots (B). Red colors represent RIN period, blue colors show noise period and black colors show silence period. Each of the six panels for A and B represent a different condition, with each condition being a variation of the periodicity of the stimulus. This MUA shows a best rate (i.e., greatest firing) of 64 Hz.

### Harmonic Complex

Example PSTHs for responsive MUA to the harmonic complex stimuli (50 trials per condition) are shown in Figure 4A, with corresponding raster plots in Figure 4B. This example shows the greatest response to the resolved harmonics of the 200 Hz missing fundamental stimulus. The peak Δ relative to baseline for this condition was 74.84 Hz when preceded by silence (with a latency of 132 ms post-stimulus onset; second panel in Figure 4), compared to a peak Δ of 59.04 Hz for gaussian noise when preceded by silence (latency of 121 ms; fifth panel in Figure 4). A total of 14/20 MUA showed pitch-selectivity using this paradigm.

**Figure 4.**
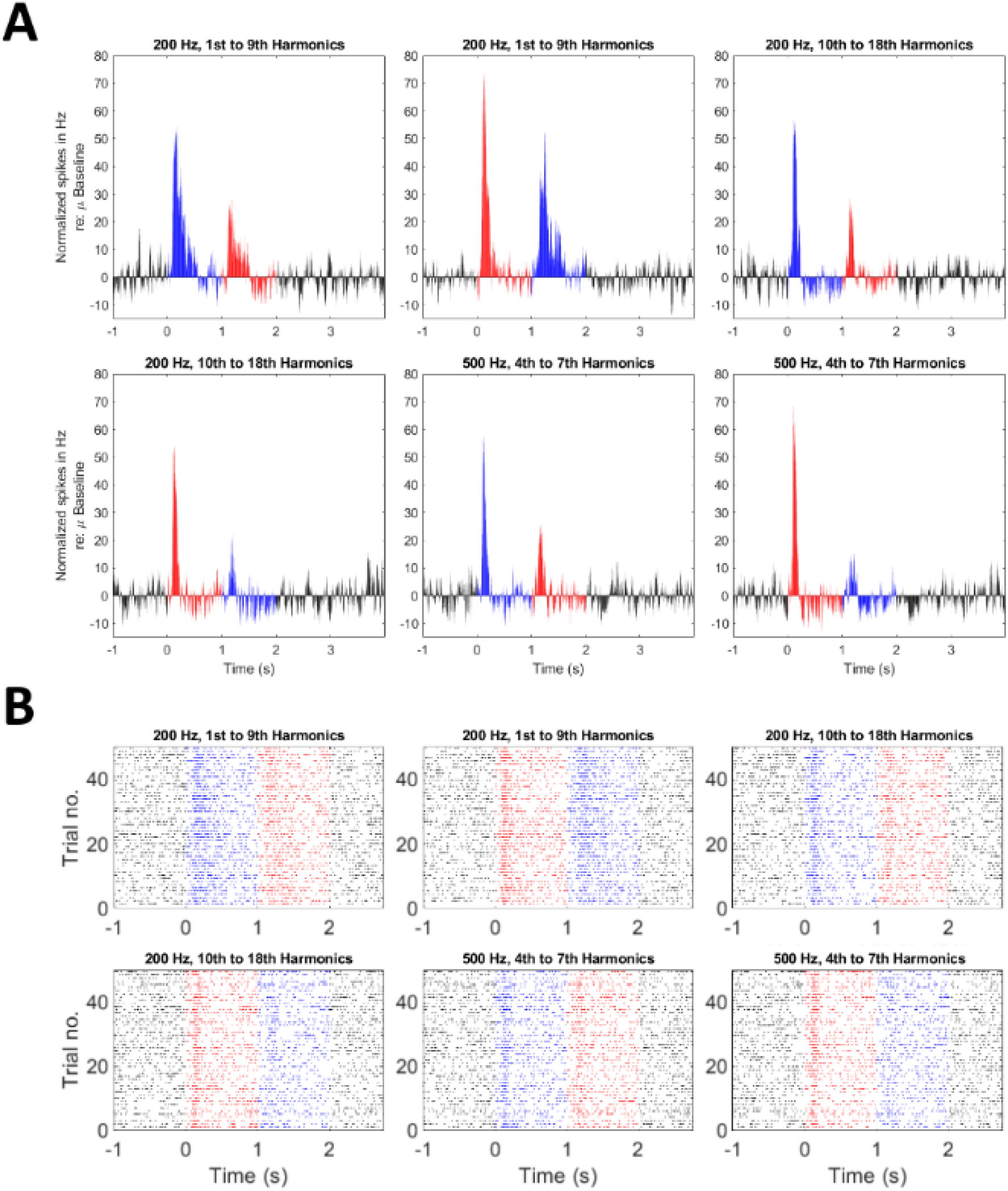
An example of normalized MUA PSTHs to the Harmonic Complex paradigm (A), with corresponding spike time raster plots (B). Red colors represent harmonic complex period, blue colors show gaussian noise period and black colors show silence period. Each of the six panels for A and B represent a different condition. From left to right, the first two panels in each subplot show responses to resolved harmonics for 200 Hz missing fundamental stimulus, the next two show higher harmonics for 200 Hz and the last two are responses to the 500 Hz missing fundamental stimulus. This MUA shows a preferential response to the lower (i.e. resolved) harmonics of the 200 Hz missing fundamental stimulus.

### Spatial distribution of pitch responsive MUA

To examine the spatial distribution of pitch and non-pitch responses, MNI coordinates of corresponding electrode contact locations were plotted on a template axial section of Heschl’s gyrus, for both pitch and non-pitch selective MUA (Figure 5). This figure shows that, regardless of the paradigm, there was no clear specific location with a greater abundance of pitch-responsive MUA, as pitch-responsive units were distributed throughout Heschl’s gyrus. Our coverage of the putative pitch region - which has been suggested to be located in anterolateral HG, mostly encompassing a region in non-primary auditory cortex and slightly extending more medially into primary auditory cortex ([19]; [20]) - was relatively limited, so we cannot rule out the possibility of a greater number/proportion of pitch-responsive units within this region. However, overall, these data are inconsistent with the concept of a singular region that is unique in representing responses related to pitch in humans.

**Figure 5.**
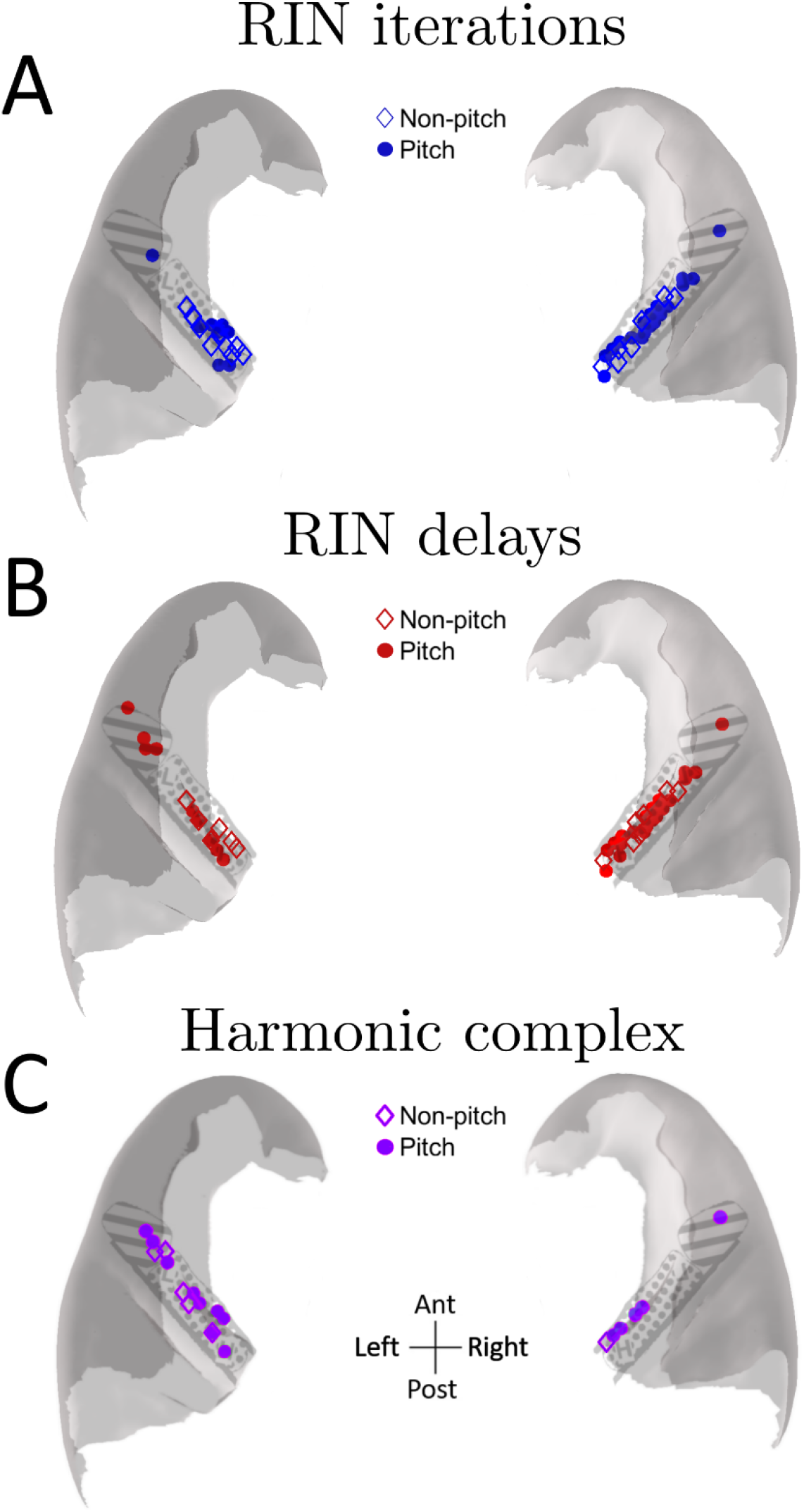
Distribution of response types for the three different paradigms: iterations (A), delays (B) and harmonic complex (C). MUA responses are plotted on a top-down, axial template section of HG, according to the MNI coordinates of their corresponding electrode contacts. Open diamonds represent non-pitch responses, filled circles indicate pitch responses. HG has been approximately sub-parcellated into posteromedial HG (dotted) and anterolateral HG (lines), based on Wallace et al. ([21]).

We also sought to determine whether there was any clear spatial distribution to the preferred periodicities of the MUA, which we termed as ‘best rate’ (Figure 6). Consistent with findings from macaques ([2]), there was no clear map of best rates, as these were distributed throughout HG. However, it should be noted that there may be some degree of sampling bias, as the majority of MUA (55.17%) responded maximally to the 256 Hz stimulus.

**Figure 6.**
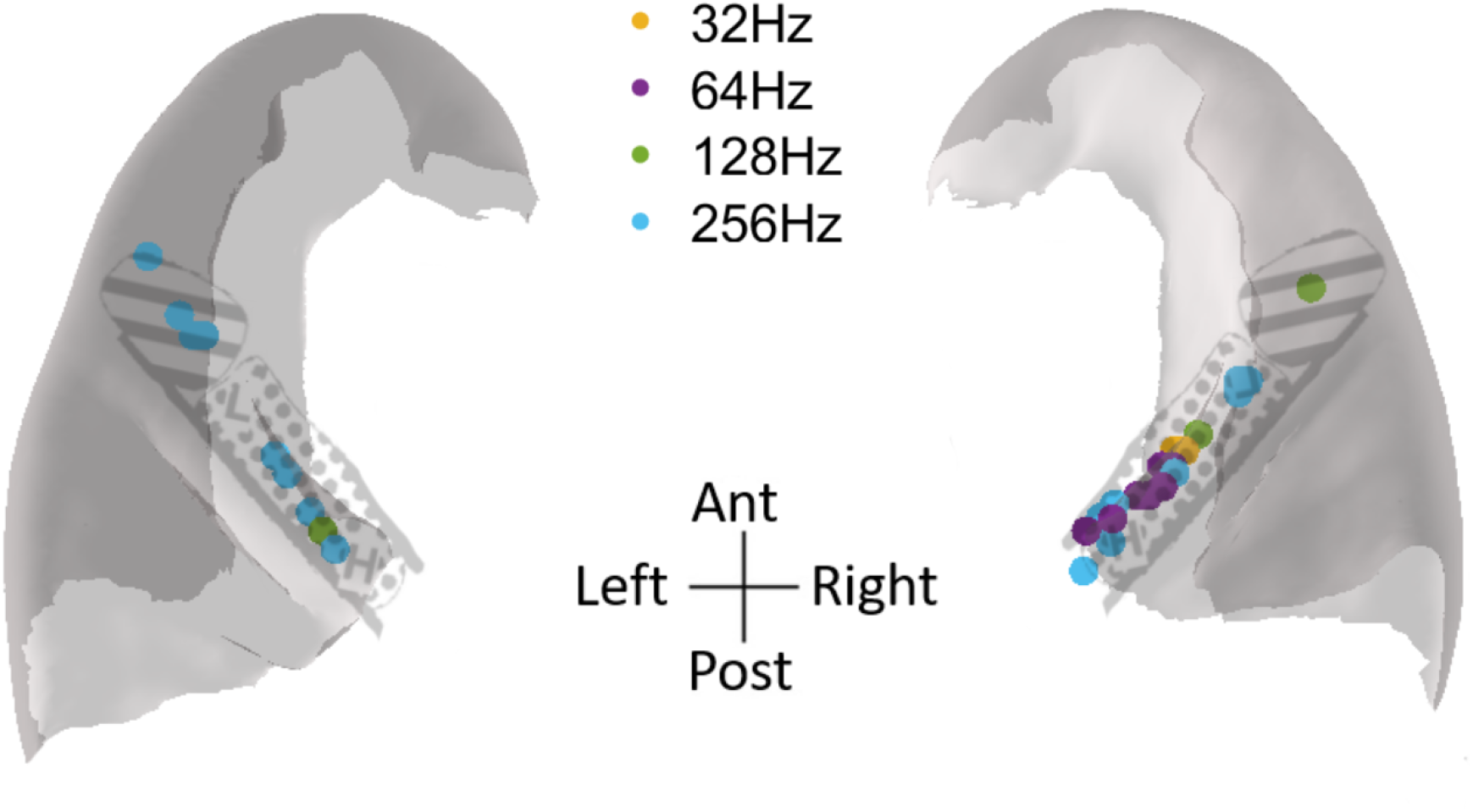
Distribution of best rates (in terms of periodicity) in response to the delays paradigm, plotted on a top-down, axial template section of HG. Sub-parcellations of posteromedial (dotted) and anterolateral HG (lines) from Wallace et al. ([21]).

## Discussion

The current study is the first of its kind in humans, wherein we have benefitted from the spatial precision of high impedance electrodes to identify the locations of multi-unit responses to pitch-like stimuli, as well as the temporal precision of electrophysiology to determine peak response latencies. These responses were distributed throughout HG, rather than confined to a particular region, thereby inconsistent with the idea of a single specialized region for pitch in humans.

### Distributed human neural responses to the pitch percept

In this study we have defined MUA associated with pitch in three different ways: 1) responses to RIN that increase with pitch salience; 2) responses to RIN that are in the range of rates associated with a pitch percept and 3) responses to harmonic complexes associated with a pitch percept. In all three paradigms we see responses that are distributed within auditory cortex and not confined to a specific region. The type of stimulus used to generate a pitch percept likely has an effect on the interpretation of results from different studies. In the current study, we attempted to mitigate this somewhat by examining responses to two different but commonly-used types of stimuli – RIN and harmonic complex stimuli. As mentioned previously, in MRI studies the type of control stimulus greatly affects interpretation of results, as the analysis often requires subtraction by design (discussed in [15] and [22]). Examining spiking activity allows for this requirement to be eased, so that we can look at more subtle changes in firing rates over time to determine responsiveness. We also were able to examine pitch specificity more robustly by reversing the presentation order of the stimuli. Therefore, we can be reasonably confident that the responses seen here are not a consequence of either the type of control stimulus used or other properties of stimulus presentation (e.g. time order effects).

Moreover, the parametric design afforded by the two RIN paradigms increased our ability to examine pitch responsiveness in greater detail – that is, by varying either the salience or the frequency of the resultant percept. In an fMRI study examining the specificity of pitch responsiveness in a region of planum temporale, it was found that the magnitude of this response did not covary with pitch salience ([23]). It is plausible that such a finding may be due to the sensitivity of the recording technique – indeed, while we did observe increasing responsiveness in pitch MUA with increasing pitch strength, across MUA these results were fairly subtle, with only ~11% change in peak firing rate between the least and most salient stimuli.

### Comparison with animal models

Animal studies have investigated pitch responses in the auditory cortex of several species, sometimes with seemingly conflicting results. In ferrets, cortical responses indicative of pitch processing have been shown to be distributed across auditory fields ([24]; [25]). Using high-field fMRI in cats, Butler et al. ([26]) found responses to RIN stimuli (compared to narrowband noise) were not present in subdivisions relating to core auditory cortex (A1 and anterior auditory field; AAF) but were instead unique to regions further upstream (posterior auditory field and A2). Bendor and Wang ([16]) demonstrated regional selectivity located in anterolateral A1 and extending slightly further medially in marmosets, while Kikuchi et al. ([2]) found responses distributed throughout macaque auditory cortex. Species differences may play a role in some disparities. Indeed, it has recently been demonstrated that there are fundamental differences in the neural representation of harmonic stimuli between humans and macaques ([27]), suggesting some degree of unique specialization in humans. With respect to this, Walker et al. ([28]) discussed across-species differences in pitch perception in the context of variations in cochlear filtering.

It is also reasonable to assume that the criteria used for the definition of pitch-responsiveness may explain discrepancies between the different studies. In Bendor and Wang ([16]), for example, a key requirement was that neurons recorded from marmosets responded similarly to harmonic complex stimuli and pure tones of a similar frequency. Due to time constraints with patients, we often did not obtain the pure tone responses on the same day for MUA in the current study, so felt that it was inappropriate to apply this criterion throughout. However, we have explored this in an additional analysis (Supplementary Figure 1) and found that contacts with pitch-responsive MUA that also responded to pure tones of a similar frequency did not spatially-localize to a single region. Likewise, this was also examined in Kikuchi et al. ([2]) in macaques, and no spatial specificity was found even when applying this criterion. Moreover, as mentioned in this latter study, it is feasible that there could be separate mechanisms for the processing of harmonic complex stimuli and pure tones, so this requirement may be overly restrictive.

The proportion of pitch responsive units here is higher than that found in macaques (~20%; [2]), although similar to proportions of voxels found in some auditory cortex regions in an imaging study utilizing RIN ([14]). In the Kikuchi et al. ([2]) study, the criteria for pitch-responsiveness were somewhat stricter than employed here, as multiple paradigms were collected in the same units. As we did not have the benefit of being able to determine that the same MUA was recorded to the different paradigms in the current study, it is possible that this factor may account for the higher proportion of units. Nonetheless, even in the previous Kikuchi et al. study, when employing stricter criteria, pitch-responses in macaques were not restricted to a single region within anterolateral auditory cortex, which is consistent with our main finding here.

### Conclusions

We have successfully demonstrated pitch responsiveness in neuronal spiking activity along the axis of Heschl’s gyrus, with sensitivity to both salience and pitch frequency. These responses were distributed throughout all regions examined, rather than restricted to a particular area. While this aspect apparently conflicts with some fMRI and marmoset studies, it is consistent with other human studies utilizing MRI ([14]) and local field potential recordings ([17]; [29]; [1]), as well as macaque ([2]) and ferret studies ([25]). As discussed in Kumar and Schönwiesner ([30]), the examination of multi-unit activity in humans is a useful step in bridging the gap between human and animal data. Future studies examining connectivity patterns between regions beyond and including HG would be beneficial in understanding the distributed nature of pitch processing, and exploring in greater detail the roles of various regions.

## Materials and Methods

### Subjects

Seven neurosurgical epilepsy patients (6 male, 1 female, all right-handed) were implanted with electrodes for the clinical purposes of identifying candidate regions of seizure foci. Sessions that were recorded within an hour of an epileptic seizure were excluded from analysis, to avoid any potential confounds. Research protocols were examined and approved by the University of Iowa Institutional Review Board.

### Electrodes and recording setup

Subjects remained in an electromagnetically-shielded facility for the duration of the recordings. Electrodes included in analyses here were a hybrid clinical-research type ([31]), with 14 exposed high-impedance contacts along the shaft, implanted along the long axis of Heschl’s gyrus (HG). All referencing was performed online, with ground and reference contacts located on the same electrode shaft. Data were recorded using a TDT-RZ2 system (Tucker-Davis Technologies), with a sampling rate of either 12207 or 24414 Hz. Precise electrode locations were confirmed by determining the MNI coordinates from electrode tracts on each individual subject’s MRI (see [1] for further details of this process).

### Experimental design and statistical analyses

#### Auditory stimuli

Stimuli were presented binaurally via Etymotic ER4B earphones coupled with custom-made earmolds. Sounds were set to a comfortable listening level for each subject. Three different stimulus paradigms were implemented here (RIN iterations, RIN delays and Harmonic Complex), all with six different conditions. The MUA was often recorded in three different paradigms on separate days or in different subjects due to time constraints, therefore an across-subject design was used to conduce analyses on the MUA recorded from the same contacts. In the RIN iterations paradigm, the first condition was a control, wherein each trial consisted of broadband noise with no iterations added. The other five of the stimulus conditions consisted of 1 second of a broadband noise period, which was either preceded or followed by RIN stimuli lasting 1.5 seconds (see [17] for a more detailed description). We randomized the order of RIN associated stimuli to control for temporal-order effects. RIN stimuli were generated using a delay-and-add algorithm with an increasing number of cycles (iterations), which results in an increasingly greater salience of the pitch percept ([32]). For this paradigm, the periodicity (i.e., pitch value) was set at either 128 or 256 Hz and was fixed for each recording. In the RIN delays paradigm, stimuli consisted of 1 second of broadband noise followed by RIN with differing periodicity (8, 16, 32, 64, 128, and 256 Hz; 1.5 seconds duration). These stimuli are useful for determining the periodicity eliciting the maximum response of MUA, defined as the best rate. RIN stimuli in both paradigms were high pass filtered at 800 Hz and normalized to the peak of the power spectral density, with broadband noise added below the 800 Hz cutoff. Finally, the third paradigm - Harmonic Complex - consisted of 1 second gaussian noise, which was either preceded or followed by a 1 second harmonic complex with the fundamental frequency missing (similar to that used in [33]). The missing fundamentals (and harmonics) for the harmonic complex were either 200 Hz (1^st^ to 9^th^ harmonics or 10^th^ to 18^th^ harmonics), or 500 Hz (4^th^ to 7^th^ harmonics). Stimuli for all paradigms were ramped (5 ms on/off) and presented with 50 repetitions per condition in a randomized order.

#### Spike sorting

Prior to spike sorting, data were down-sampled offline to 12kHz and denoised using the demodulated band transform ([34]). In order to threshold multi-unit data from noise, spike sorting was performed for each recording block using the MountainSort algorithm ([35]), which offers a fast and fully automated approach to spike sorting, utilizing both principal component analysis and the ISO-SPLIT clustering algorithm ([36]). Briefly, data were first bandpass filtered between 300 and 6000 Hz, whitened to decorrelate the data across channels, and then run through the MountainSort algorithm. Sorted spikes were imported into Matlab for visualization and manual curation. For example, separate clusters of MUA were combined on each electrode (as extracted spike waveforms did not conform to a classical depiction of well-isolated single units with low variability), and any spuriously high amplitudes (> 500 μV) were removed before epoching. All spike sorted data were then epoched around the stimulus for each condition, beginning 1 second before first stimulus onset to 4 seconds after.

#### Analyses

Peri-stimulus time histograms (PSTHs) were produced for each auditory stimulus condition. Auditory responsive channels were defined as increasing their firing rate to the auditory stimulus by greater than 3 standard deviations above the mean baseline firing rate (−1000 to −100 prestimulus) for a minimum of 25 ms at the onset of the stimulus. Responsiveness to pitch-like stimuli was determined using the same criteria. Latencies were extracted from averaged PSTHs as the timing of the maximum response. For the RIN iterations paradigm, MUA was classified as pitch-responsive if it showed a selective increase to increasing pitch salience that was also evident in conditions where the transition was reversed (i.e. RIN to noise), in order to account for any time order or pop-out effects of the RIN stimulus. Pitch selectivity for the RIN delays paradigm was defined as preferential responsiveness that was evident above the lower limit of pitch (~30 Hz; [9]), while best rates were determined to the same stimuli as the periodicity of the RIN that evoked the greatest increase in firing within the first 500 ms of the RIN stimulus onset. For the harmonic complex paradigm, pitch MUA was defined as MUA that responded preferentially to a particular harmonic complex stimulus. Channel locations were plotted on an MNI template of HG, with each of their MNI coordinates used to indicate their location along the axis of HG, and different symbols used to indicate whether MUA showed selective responses to pitch-like stimuli or no selectivity at that location. For display purposes, whenever a contact had multiple clusters of MUA (i.e. across different sessions), x-coordinates were shifted by 1mm to not overlap – in the supplementary Excel file of MNI coordinates, these were left intact. Differences in proportions of pitch/non-pitch MUA for the three different paradigms were tested for significance with multiple Fisher’s exact tests in GraphPad Prism version 9 for Windows (GraphPad Software, San Diego, California USA).

## Acknowledgements

This work was supported by a Wellcome Trust grant (WT106964MA) and a Medical Research Council grant (MR-T032553-1) awarded to Timothy D Griffiths and an NIH grant (DC004290) awarded to Matthew A Howard. We thank Haiming Chen, Rachel Gold and Richard Reale for their help with conducting the experiments, and Christopher Garcia with assistance in collating subject information.

---

**Supplementary Figure 1.**
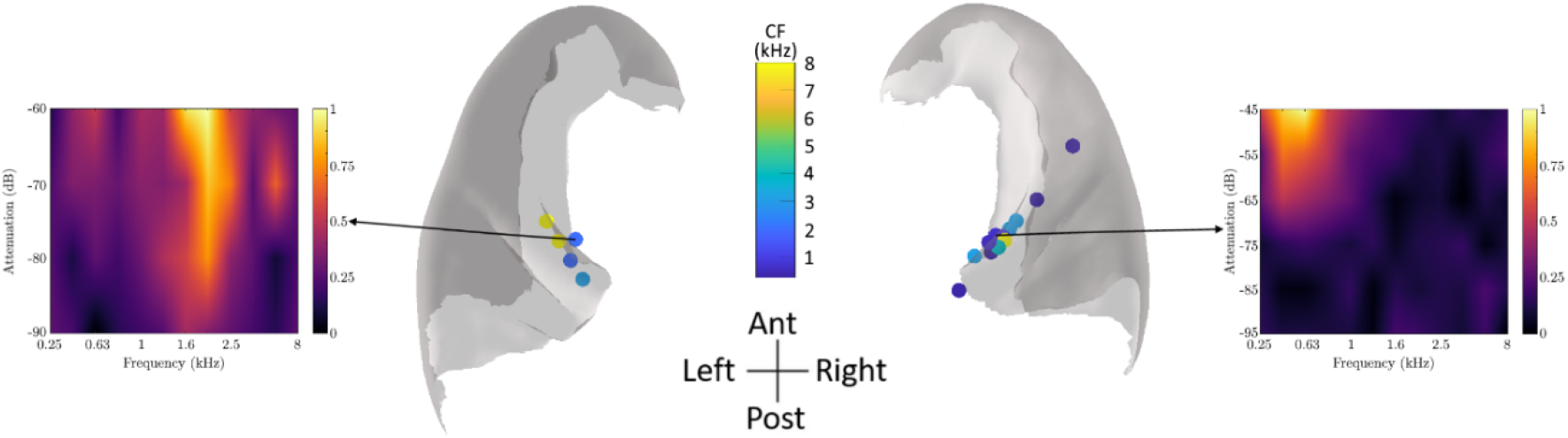
Center figure shows distribution of pitch-responsive contacts across all paradigms, wherein MUA also showed a significant response to pure tones of a similar frequency (within 1 octave). Contacts are color-coded according to their characteristic frequency (CF) determined by frequency response areas and PSTHs, constructed from responses to tones of varying frequency and amplitude (n = 17 unique contacts). Examples of these frequency response areas – normalized according to maximum firing rate – are shown either side. Both of these examples showed responses to pure tones of similar frequencies to the pitch-eliciting stimuli, though one had a higher CF (left) and had a lower CF (right). Note that a number of contacts had MUA recorded from them across different sessions in response to the pitch stimulus paradigms.

## Notes

### Competing Interest Statement

The authors have declared no competing interest.

